# Genomic interrogation of the burden and transmission of multidrug-resistant pathogens within and across hospital networks

**DOI:** 10.1101/764787

**Authors:** Norelle L. Sherry, Robyn S. Lee, Claire L. Gorrie, Jason C. Kwong, Rhonda L. Stuart, Tony Korman, Caroline Marshall, Charlie Higgs, Hiu Tat Chan, Maryza Graham, Paul D.R. Johnson, Marcel Leroi, Caroline Reed, Michael Richards, Monica A. Slavin, Leon J. Worth, Benjamin P. Howden, M. Lindsay Grayson, on behalf of the Controlling Superbugs Study Group

**Author notes:** Corresponding author: Prof. M. Lindsay Grayson, Department of Infectious Diseases, Austin Health., and Alternate corresponding author: Prof. Benjamin Howden, MDU Public Health Laboratory, Department of Microbiology & Immunology at the Peter Doherty Institute for Infection and Immunity, University of Melbourne. Current affliliation: Epidemiology Division, Dalla Lana School of Public Health, University of Toronto, Toronto, Canada, and Centre for Communicable Disease Dynamics, Harvard T.H. Chan School of Public Health. Authors contributed equally (Benjamin P. Howden, M. Lindsay Grayson). Key points: We conducted a prospective multi-center study to investigate multidrug-resistant organisms (MDROs), using genomics to define the burden, distribution, and transmission of MDROs within and between hospital networks, defining a new benchmarking measure for future genomic studies.

## Abstract

**Background:** Multidrug-resistant organisms (MDROs) disproportionately affect hospitalized patients due to the combination of comorbidities, frequent antimicrobial use, and in-hospital MDRO transmission. Identification of MDRO transmission by hospital microbiology laboratories is difficult due to limitations of existing typing methods.

**Methods:** We conducted a prospective multicenter genomics implementation study (8 hospitals, 2800 beds) from 24^th^ April to 18^th^ June 2017 in Melbourne, Australia. Clinical and screening isolates from hospital inpatients were collected for six MDROs (*vanA* VRE, MRSA, ESBL *E. coli* [ESBL-Ec] and *Klebsiella pneumoniae* [ESBL-Kp], and carbapenem-resistant *Pseudomonas aeruginosa* [CRPa] and *Acinetobacter baumannii* [CRAb]), sequenced (Illumina NextSeq) and analyzed using open-source tools. MDRO transmission was assessed by genomics (core SNP phylogeny, grouped by species and ST) and compared to epidemiologic data.

**Results:** 408 isolates were collected from 358 patients; 47.5% were screening isolates. ESBL-Ec was most common (52.5%), then MRSA (21.6%), *vanA* VRE (15.7%) and ESBL-Kp (7.6%).

We define the transmission rate for each MDRO by genomics and epidemiology; 31.6% of all study patients had potential genomic links to other study isolates; 86% of these were confirmed by epidemiologic links (probable or possible transmission). The highest transmission rates occurred with *van*A VRE (88.4% of patients).

**Conclusions:** Combining genomics with high-quality epidemiologic data gives substantial insights into the burden and distribution of critical MDROs in hospitals, including in-hospital transmission. By defining transmission rates by genomics, we hope to enable comparisons over time and between sites, and introduce this as a new outcome measure to assess the efficacy of infection control interventions.

## Introduction

Multidrug-resistant organisms (MDROs) are increasing globally, and disproportionately affect hospital patients [1, 2]. Infections with these pathogens may be acquired in healthcare settings or in the community, and are associated with increased morbidity, mortality, length of hospital stay and healthcare costs [2-4]. Whilst many healthcare systems, including those in Australia, have successfully implemented surveillance programs for low-burden, high-impact pathogens, such as carbapenemase-producing Enterobacteriaceae (CPE)[5-10], these surveillance systems do not always comprehensively address more common MDROs, resulting in incomplete data about some MDROs frequently affecting patients, such as extended-spectrum beta-lactamase-producing *E. coli* (ESBL-Ec) or methicillin-resistant *Staphylococcus aureus* (MRSA).

In Australia, as in Europe, the major pathogen surveillance programs such as Australian Group for Antimicrobial Resistance (AGAR) and the Victorian Hospital Pathogens Surveillance System (VHPSS) [11, 12] focus on isolates from bacteremic patients, meaning the true burden of MDROs, many of which lead to colonization or non-bacteremic infection, is undefined and poorly understood [9, 11]. Australia has one the highest rates of VRE in the world (47.0% in 2017 [13]), having been dominated by *vanB* until the last five years, when *vanA* VRE has emerged rapidly across multiple states [14]. Whilst the patient risk factors of MDRO acquisition are well understood for most of these organisms [15-22], further analysis using whole genome sequencing has the potential to add further insights, particularly defining the relatedness of isolates by genomics (and hence putative transmission when combined with epidemiologic data), which to date has only just started to be applied in a clinical setting [23, 24].

In this genomics implementation and evaluation study, we performed comprehensive surveillance of the clinical and genomic epidemiology of MDROs across multiple hospitals over a two-month period, to (i) estimate the local burden of MDRO infection and colonization, (ii) assess risk factors for MDRO acquisition, (iii) establish the local population structure and compare to national and international data, and (iv) investigate the use of genomics to predict in-hospital MDRO transmission, and define a transmission rate per occupied bed days.

## Methods

An overview of methods (including inclusion and exclusion criteria) is given in Figure 1, with more detailed information available in Supplementary Data.

**Figure 1.**
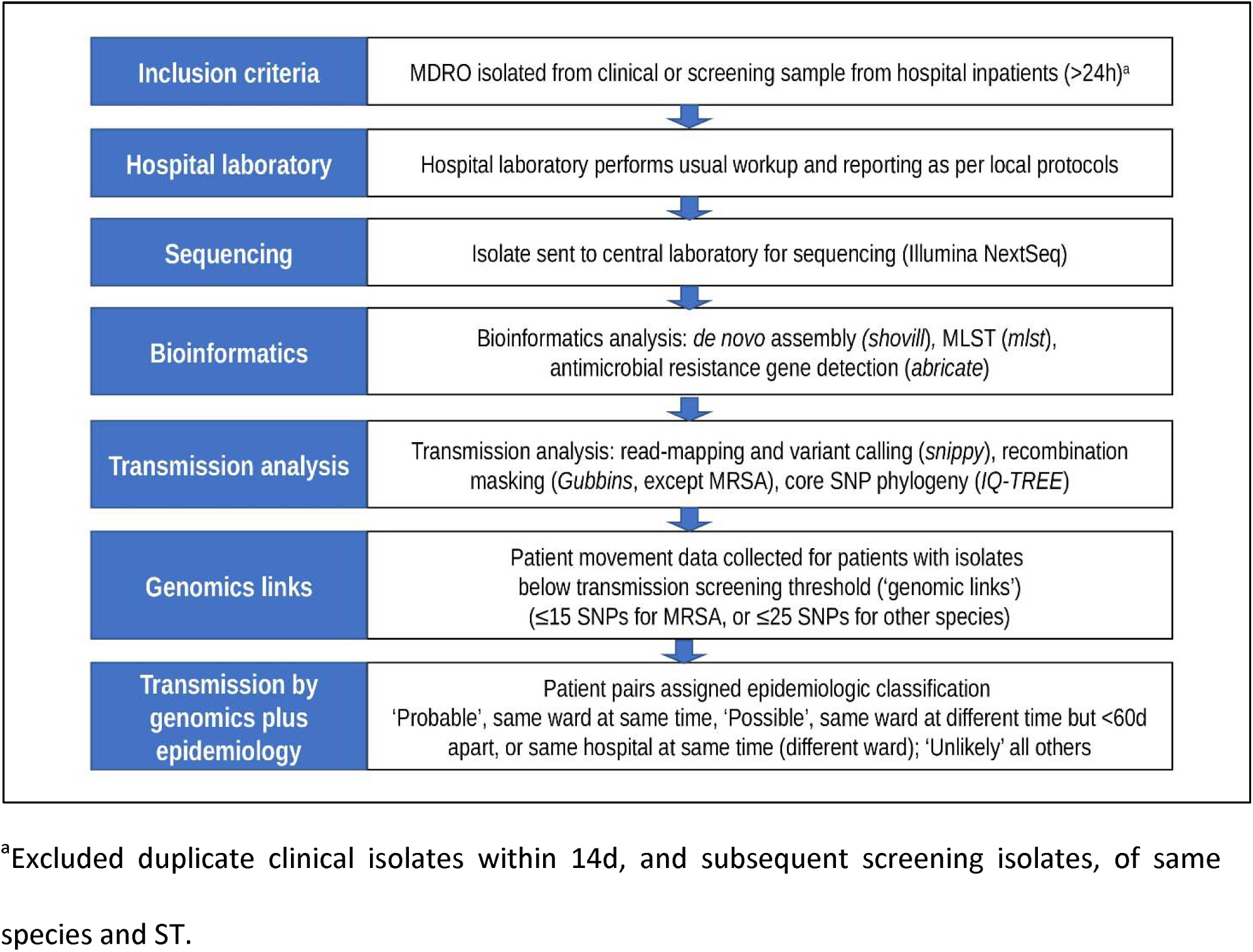
Overview of study design. Note: carbapenemase-producing Enterobacterales excluded as already covered by existing state-wide genomic surveillance program [5]. MLST, multilocus sequence typing; SNP, single nucleotide polymorphism. Names of bioinformatics tools listed in italics. References for bioinformatics tools: *shovill* (v1.0.4; https://github.com/tseemann/shovill), *mlst* (https://github.com/tseemann/mlst), *abricate* (v0.9.5, https://github.com/tseemann/abricate), *snippy* (https://github.com/tseemann/snippy), *Gubbins* [25], IQ-TREE [26].

### Study design

We conducted a prospective multicenter study of eight hospital sites from four hospital networks (Table 1), covering approximately 2800 acute and subacute (aged care/rehabilitation) patient beds. Isolates were collected during an eight-week pilot study (24^th^ April to 18^th^ June 2017), conducted as part of a larger study for the Melbourne Genomics Health Alliance, using genomics for MDRO surveillance in hospitals. Clinical and screening isolates of six MDROs were collected from hospital inpatients: *vanA* vancomycin-resistant *Enterococcus* faecium (*vanA* VRE), methicillin-resistant *Staphylococcus aureus*, extended-spectrum beta-lactamase (ESBL)-phenotype *Escherichia coli* and *Klebsiella pneumoniae* (ESBL-Ec and ESBL-Kp), carbapenem-resistant *Acinetobacter baumannii* complex (CRAb) and carbapenem-resistant *Pseudomonas aeruginosa* (CRPa)(Table 2). Carbapenem-resistant Enterobacterales (CPE) were excluded, as these were already collected for a comprehensive state-wide CPE surveillance program [5, 27]. Whilst vanB VRE are dominant in Australia, we elected to focus on vanA VRE as it emerged more recently in Victoria, has a greater level of associated antimicrobial resistance and costs, and could potentially be more amenable to infection control interventions.

**Table 1.** Hospital sites and characteristics.

| Hospital network | Hospital code | Hospital description | No. of inpatient beds | High-risk wards | MDRO screening practices during study period |
| --- | --- | --- | --- | --- | --- |
| **A** | **A1** | Tertiary referral center, including ICU, solid organ and bone marrow transplant | 560 | ICU<br>Hematology/BMT and Oncology<br>Renal Transplant<br>Liver Transplant | ICU, hematology/oncology, renal and liver transplant wards screened on admission and twice weekly for *vanA* VRE and MRGN<br>Additional MRSA screening in ICU (on admission and twice weekly)<br>Quarterly point-prevalence survey for *vanA* VRE and MRGN<br>MRSA screening before critical surgeries (prosthetic joint, spinal and cardiac) |
|  | **A2** | Subacute hospital, aged care and rehabilitation services | 150 | None | Quarterly point-prevalence survey for *vanA* VRE and MRGN |
|  | **A3** | Subacute hospital, rehabilitation services | 60 | None | Quarterly point-prevalence survey for *vanA* VRE and MRGN |
| **B** | **B1** | Tertiary referral center, including ICU and solid organ transplant and specialist pediatric hospital (including neonatal ICU) | 640 | ICU<br>Renal Transplant | ICU and renal ward screened for *vanA* VRE and carbapenem-resistant Gram negatives (CRGN) on admission and weekly<br>MRSA screening before cardiac surgery |
|  | **B2** | Tertiary referral center, including ICU and some aged care & rehabilitation services | 573 | ICU | ICU patients screened for *vanA* VRE and carbapenem-resistant Gram negatives (CRGN) on admission and weekly |
| **C** | **C1** | Tertiary referral center, including ICU, solid organ and bone marrow transplant | 571 | ICU<br>Hematology/BMT | ICU and hematology ward screened on admission and weekly for *vanA* VRE and MRGN |
|  | **C2** | Subacute hospital, aged care and rehabilitation services | 150 | None | None |
| **D** | **D1** | Specialized cancer care center. Located adjacent to Hospital 3A (ICU patients cared for at 3A before transfer back to hospital 4) | 96 | Hematology | Hematology ward patients screened on admission and weekly for *vanA* VRE and MRGN |

ICU, intensive care unit; MRGN, multi-resistant Gram negatives (includes ESBL and carbapenem-resistant phenotypes); BMT, bone marrow transplant (allogeneic).

**Table 2.** Laboratory definitions of MDROs.

| Organism | Laboratory definition |
| --- | --- |
| *vanA* vancomycin-resistant<br>*Enterococcus faecium* (*vanA* VRE) | Positive *vanA* PCR result (including *vanA+B*) |
| Methicillin-resistant *Staphylococcus aureus* (MRSA) | Positive cefoxitin screen (disc <22mm or MIC>4mg/L) or oxacillin MIC >2mg/L |
| Ceftriaxone non-susceptible *E. coli*<br>or *K. pneumoniae* (ESBL-Ec/ESBL-Kp) | *E. coli* or *K. pneumoniae* with ceftriaxone MIC ≥4mg/L<br>Excludes carbapenemase-producing Enterobacteriaceae (collected under statewide CPE surveillance program). AmpC phenotypes included. |
| Carbapenem non-susceptible<br>*P. aeruginosa* (CRPa) | Meropenem MIC ≥8mg/L (resistant by CLSI criteria [28], intermediate/resistant by EUCAST criteria [29]) AND resistant to piperacillin-tazobactam AND ceftazidime |
| Carbapenem non-susceptible<br>*A. baumannii* (CRAb) | Meropenem MIC ≥8mg/L (resistant by CLSI criteria [28], intermediate/resistant by EUCAST criteria [29]) |

### MDRO screening protocols

Existing MDRO screening protocols varied between hospitals (Table 1); further details are available in Supplementary Data. Hospital infection control practices (including patient isolation and organism-specific terminal cleaning practices) were assessed at baseline and at the conclusion of the study; no changes were made during the study period. Results of genomic analyses were not available to hospitals during the study period.

### Sequencing laboratory workflow and bioinformatics analysis

See Figure 1 and Supplementary Data for detailed methods (including Supplementary Table S1 [References genomes used for transmission analysis]).

This study was approved by the Melbourne Health Human Research Ethics Committee (HREC) and endorsed by the corresponding HREC at each participating site.

### Data availability

Raw sequence data has been uploaded to the Sequence Read Archive under BioProject PRJNA****** (to be uploaded soon).

## Results

### Isolate numbers, patient demographics and specimen types

During the eight-week study period (24^th^ April to 18 ^th^ June 2017), 408 MDRO isolates from 358 patients were collected; most patients (88.3%) only had a single isolate included, 10.1% had two isolates, and 1.7% had three or more isolates. The median age of patients was 67yo, and was similar for most species except CRAb (only two patients)(Table 3).

**Table 3.** Summary of patient and isolate numbers.

|  | Species |  |  |  |  |  | Overall |
| --- | --- | --- | --- | --- | --- | --- | --- |
|  | ESBL-Ec | MRSA | *vanA* VRE | ESBL-Kp | CRPa | CRAb |  |
| No. isolates (% total) | 214 (52.5%) | 88 (21.6%) | 64 (15.7%) | 31 (7.6%) | 9 (2.2%) | 2 (0.5%) | 408 |
| No. patients (% total)<sup>a</sup> | 203 (56.7%) | 86 (24.0%) | 60 (16.8%) | 30 (8.4%) | 8 (2.2%) | 2 (0.6%) | 358 |
| % Male (% total patients)<sup>a</sup> | 51.7% | 54.7% | 61.7% | 66.7% | 62.5% | 50.0% | 55.3% |
| Age in yrs (median, range) | 68 (1-100) | 62 (1-97) | 67 (26-93) | 66.5 (20-89) | 65.5 (28-82) | 52.5 (29-76) | 67 (1-100) |

Abbreviations: ESBL-Ec, extended-spectrum beta-lactamase phenotype *E. coli*; MRSA, methicillin-resistant *S. aureus*; *vanA* VRE, *vanA*-producing vancomycin-resistant *E. faecium*; ESBL-Kp, extended-spectrum beta-lactamase phenotype *K. pneumoniae*; CRPa, carbapenem-resistant *P. aeruginosa* (also resistant to piperacillin-tazobactam and ceftazidime); CRAb, carbapenem-resistant *A. baumannii*.
<sup>a</sup> Twenty-eight patients had more than one species isolated, hence percentages add to >100%

Overall, 47.5% of isolates were collected for screening purposes, although this varied between species; the majority of *vanA* VRE isolates were from screening samples (81.3%), whereas the majority of MRSA isolates (92.0%) were from clinical samples (collected for suspected infection)(Figure 2a and Supplementary Table S2). Of the clinical samples, urine specimens were most common (45.8% of clinical isolates), followed by non-sterile sites (25.2%), blood cultures (10.7%), respiratory specimens (9.8%) and other sterile sites (8.4%). Blood cultures represented a small proportion of clinical isolates for most species (8.6% [MRSA] to 14.3% [ESBL-Kp]), except *vanA* VRE (33.3% of clinical isolates, although small number of clinical isolates overall [n=12]).

**Figure 2.**
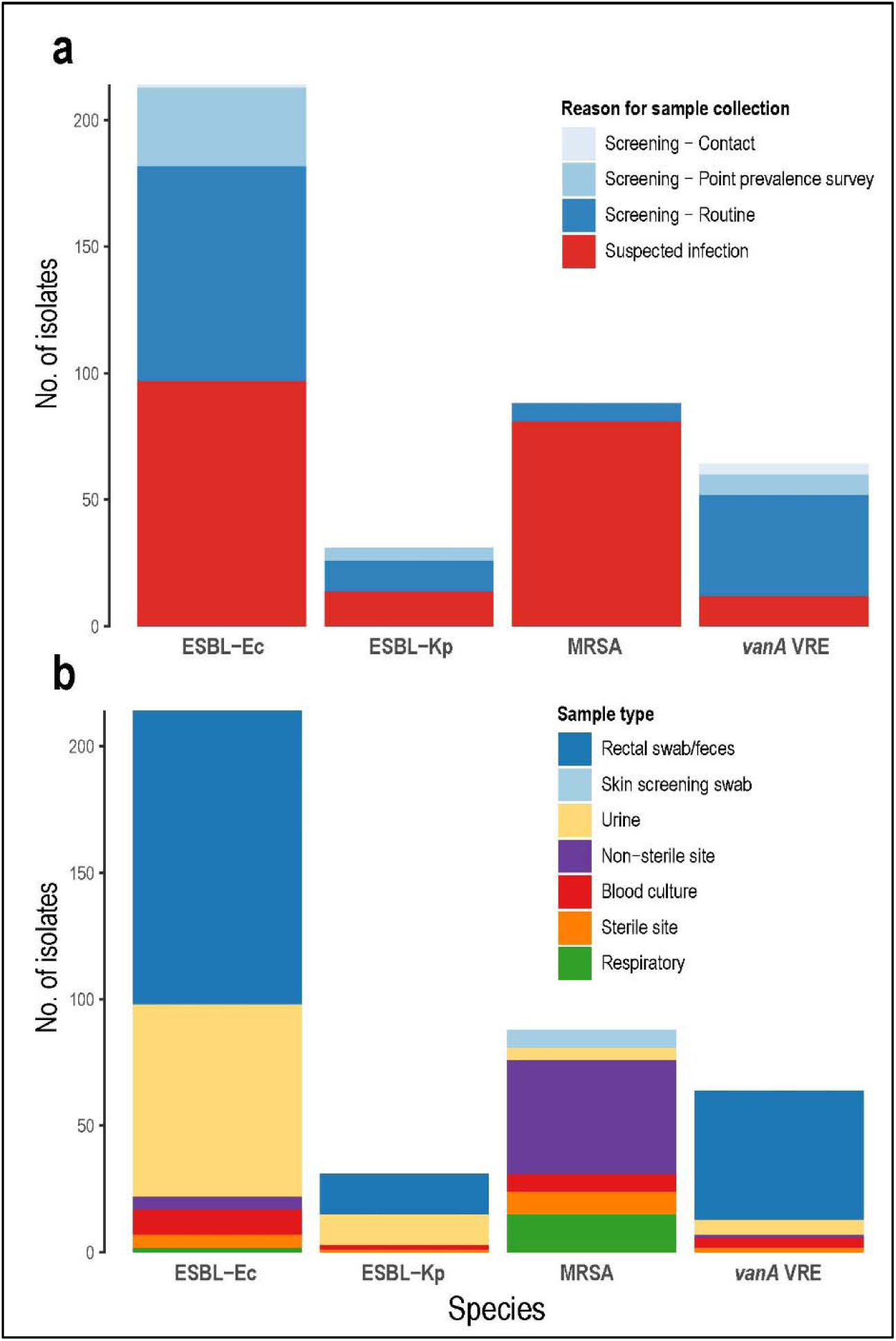
Characteristics of isolates - reason for sample collection, and specimen type. **Fig 2a** Reason for sample collection; **Fig 2b** Sample type Abbreviations: ESBL-Ec, extended-spectrum beta-lactamase phenotype *E. coli*; MRSA, methicillin-resistant *S. aureus*; *vanA* VRE, *vanA*-producing vancomycin-resistant *E. faecium*; ESBL-Kp, extended-spectrum beta-lactamase phenotype *K. pneumoniae*. See Supplementary Tables S2 and S3 for further details.

### High rates of MDRO isolation, especially ESBL-Ec and MRSA

To define the incidence of each MDRO (and to enable comparisons between different wards and hospital sites), we calculated rates per 100,000 occupied bed days. MDRO infections occurred at a rate of 107.1 patients per 100,000 occupied bed days (OBDs), whilst the overall MDRO burden (both infection and colonization) was 294.5 patients per 100,000 OBDs. Considering infection only (as this not affected by different screening practices), rates were much higher in patients on high-risk wards or ICU (infection rate, 151.1, and total burden, 900.4 patients per 100,000 OBDs)(Figure 4 and Table 4). ESBL-Ec infections were most common, followed by MRSA and ESBL-Kp, whilst colonization rates were highest in ESBL-Ec and *vanA* VRE. CRPa and CRAb were uncommon in participating sites (total burden 6.3 and 1.6 patients per 100,000 OBDs respectively).

**Table 4.** Rates of patient MDRO infection and/or colonization per 100,000 occupied bed days^a^.

|  | Species |  |  |  |  |  | Overall |
| --- | --- | --- | --- | --- | --- | --- | --- |
|  | ESBL-Ec | MRSA | vanA VRE | ESBL-Kp | CRPa | CRAb |  |
| **All wards** |  |  |  |  |  |  |  |
| MDRO infections | 50.0 | 42.9 | 4.8 | 6.3 | 3.2 | 0.0 | 107.1 |
| MDRO colonization | 112.7 | 25.4 | 42.9 | 17.5 | 3.2 | 1.6 | 203.2 |
| Total burden | 152.4 | 66.7 | 44.4 | 23.0 | 6.3 | 1.6 | 294.5 |
| Clinical isolates | 77.0 | 64.3 | 9.5 | 11.1 | 6.3 | 1.6 | 169.8 |
| Blood cultures | 7.9 | 5.6 | 3.2 | 1.6 | - | - | 18.3 |
| **High-risk wards & ICU<sup>b</sup>** |  |  |  |  |  |  |  |
| MDRO infections | 50.4 | 63.0 | 18.9 | 6.3 | 12.6 | 0.0 | 151.1 |
| MDRO colonization | 421.9 | 50.4 | 226.7 | 37.8 | 25.2 | 12.6 | 774.5 |
| Total burden | 453.3 | 107.0 | 245.6 | 44.1 | 37.8 | 12.6 | 900.4 |
| **MDRO infection** |  |  |  |  |  |  |  |
| Relative risk for high-risk wards compared to other wards | 0.97 | 1.54 | 6.93 | 0.99 | 6.93 | - | 1.46 |

Abbreviations: ESBL-Ec, extended-spectrum beta-lactamase *E. coli*; MRSA, methicillin-resistant *S. aureus*; vanA VRE, vanA-producing vancomycin-resistant *E. faecium*; ESBL-Kp, extended-spectrum beta-lactamase *K. pneumoniae*; CRPa, carbapenem-resistant *P. aeruginosa* (also resistant to piperacillin-tazobactam and ceftazidime); CRAb, carbapenem-resistant *A. baumannii*; MDRO, multidrug-resistant organism.
<sup>a</sup> Occupied bed days - number of patients admitted overnight (excluding mental health and hospital-in-the-home services).
<sup>b</sup> High-risk wards, includes hematology, oncology, renal ward (including renal transplant), and liver transplant wards; ICU, intensive care unit.
Note: Total burden less than infection + colonization as duplicates excluded.

**Figure 3.**
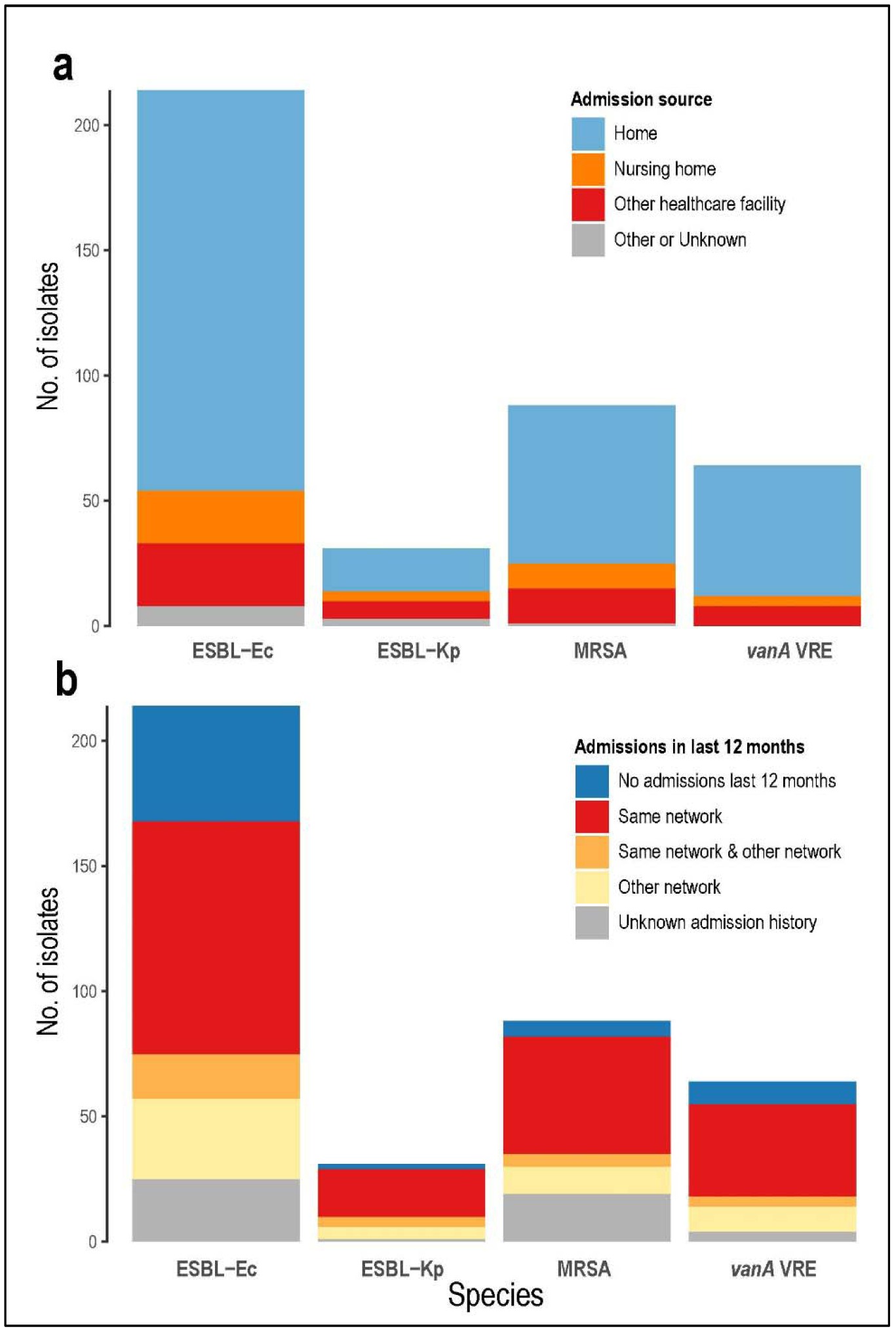
Patient admission source and history. **Fig 3a** Admission source (where patient was admitted from); **Fig 3b** Admission history in previous 12 months. See Supplementary Table S2 for further details

**Figure 4.**
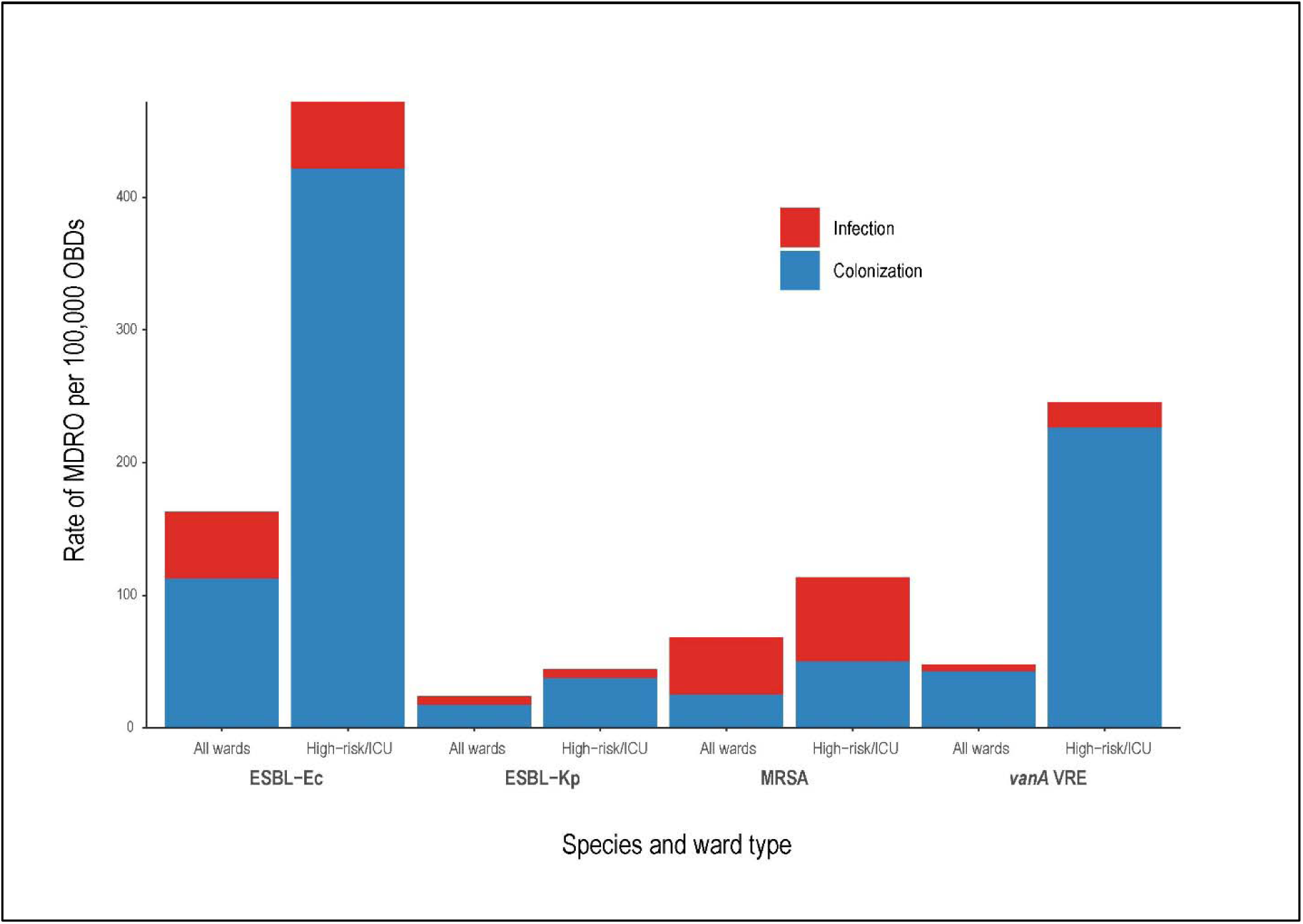
Rates of MDRO infection and colonization per 100,000 occupied bed days (OBDs) Abbreviations: ESBL-Ec, extended-spectrum beta-lactamase phenotype *E. coli*; MRSA, methicillin-resistant *S. aureus*; *vanA* VRE, *vanA*-producing vancomycin-resistant *E. faecium*; ESBL-Kp, extended-spectrum beta-lactamase phenotype *K. pneumoniae*. High-risk wards include hematology, oncology, renal ward (including renal transplant), liver transplant ward, and ICU (intensive care unit). Occupied bed day defined as number of beds occupied by patients at midnight, excluding day cases, mental health and hospital-in-the-home. See Supplementary Table S4 for more detailed data

### There are differences in time-to-collection profiles between MDROs

To investigate the likely location of MDRO acquisition, 193/333 (58.0%) non-duplicate isolates were collected within the first two days of hospital admission (excluding patients transferred from other hospitals). The majority of MDROs were isolated within the first week of admission (0-7 days, excluding patients transferred from other healthcare facilities), particularly MRSA (86.3%), ESBL-Ec (81.0%) and ESBL-Kp (70.8%). In contrast, only 60.0% of CRPa and 50% of *vanA* VRE were collected in the first week of admission. The two CRAb isolates were both from clinical samples taken in the second week of admission from patients transferred from overseas. These proportions were similar for both screening and clinical isolates, except for *vanA* VRE, where more clinical isolates were detected in weeks two and three of admission (25% and 33% of *vanA* VRE screening isolates, respectively), compared to week one (Figure 5).

**Figure 5.**
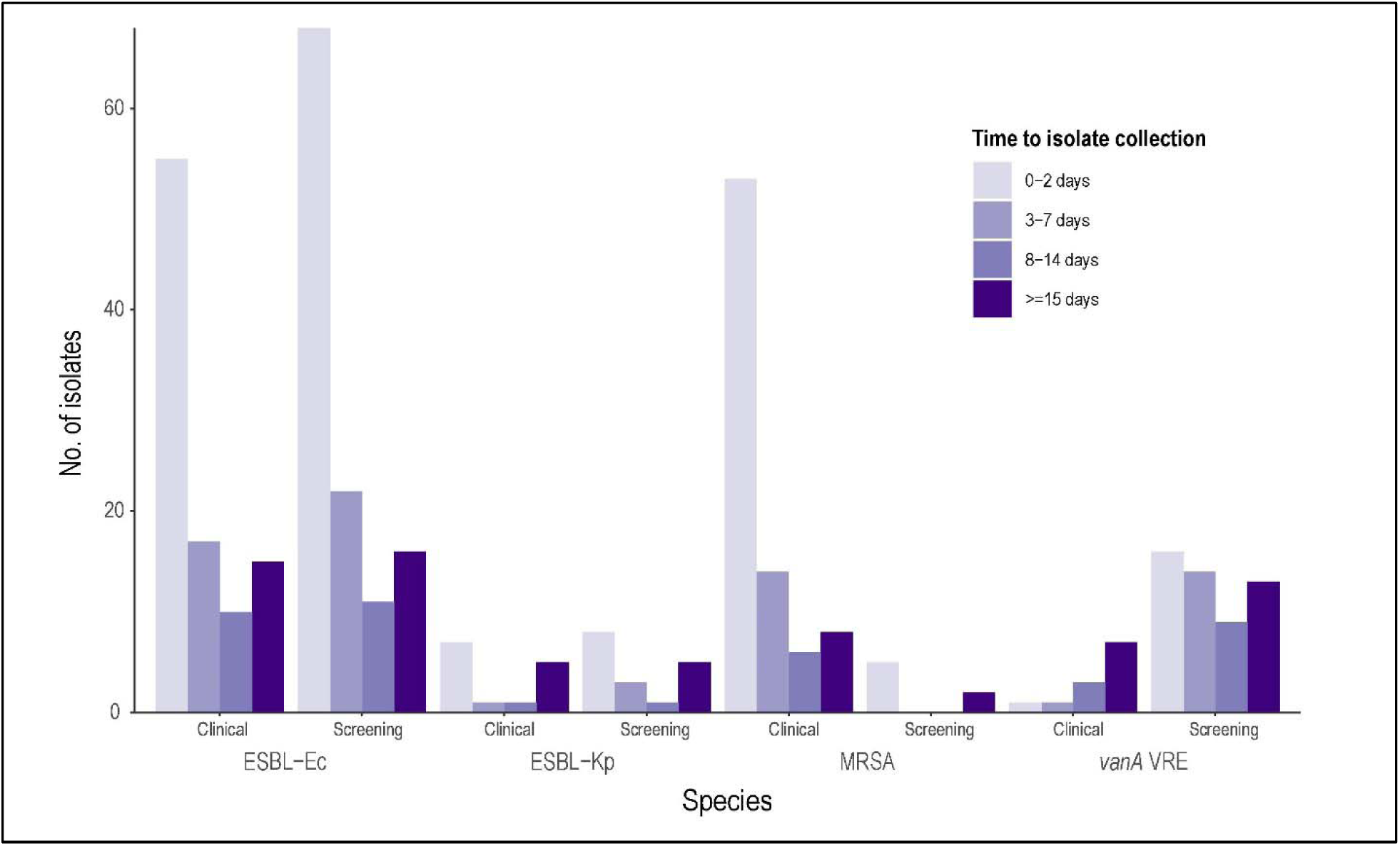
Time from patient admission to isolate collection, by species and reason for sample collection. Abbreviations: ESBL-Ec, extended-spectrum beta-lactamase phenotype *E. coli*; MRSA, methicillin-resistant *S. aureus*; *vanA* VRE, *vanA*-producing vancomycin-resistant E. *faecium*; ESBL-Kp, extended-spectrum beta-lactamase phenotype *K. pneumoniae*. ^a^Percentage of clinical or screening isolates of this species Clinical isolates, from samples collected for suspected infection. Screening isolates, from samples collected for MDRO surveillance.

### Very few MDROs were isolated from patients without healthcare contact

25.7% of MDROs were isolated from patients with a known history of colonization with that MDRO in the previous 12 months; 60.7% of these were thought to represent infection rather than colonization (mostly ESBL-Ec and MRSA)(data not shown).

Only a small number of patients (42 patients, 11.7%) had MDROs isolated without a history of healthcare exposure (admitted from home, no known admissions in last 12m, not known to be colonized in last 12m or unknown colonization status)(Figure 3). Most of these were ESBL-Ec (32 patients, 25.0% were clinical isolates); 18/32 patients had ESBL-Ec isolates within the first two days of admission. In contrast, only four patients with MRSA (4.6% of MRSA), and six patients with *vanA* VRE (10.0% of VRE) had a similar lack of healthcare exposure (as defined above). Further data regarding wards and medical units where MDROs were isolated are detailed in Supplementary Table S2.

### Molecular epidemiology reveals unique features of local MDRO population structures

To investigate the local population structures of each MDRO and allow comparisons to national and international data, we examined the most common multilocus sequence types for each species. Population structures figures by eBURST methodology for all species are available in Supplementary Figure 1.

#### vanA VRE

Of the *vanA* VRE isolates, ST1421 was the dominant sequence type (62.5%), followed by ST203 (28.1%)(Figure 6). Notably, ST1424 (a dominant ST in neighbouring states of Australia) and ST 796 (a common ST amongst Australian *vanB* isolates) were absent [13].

**Figure 6.**
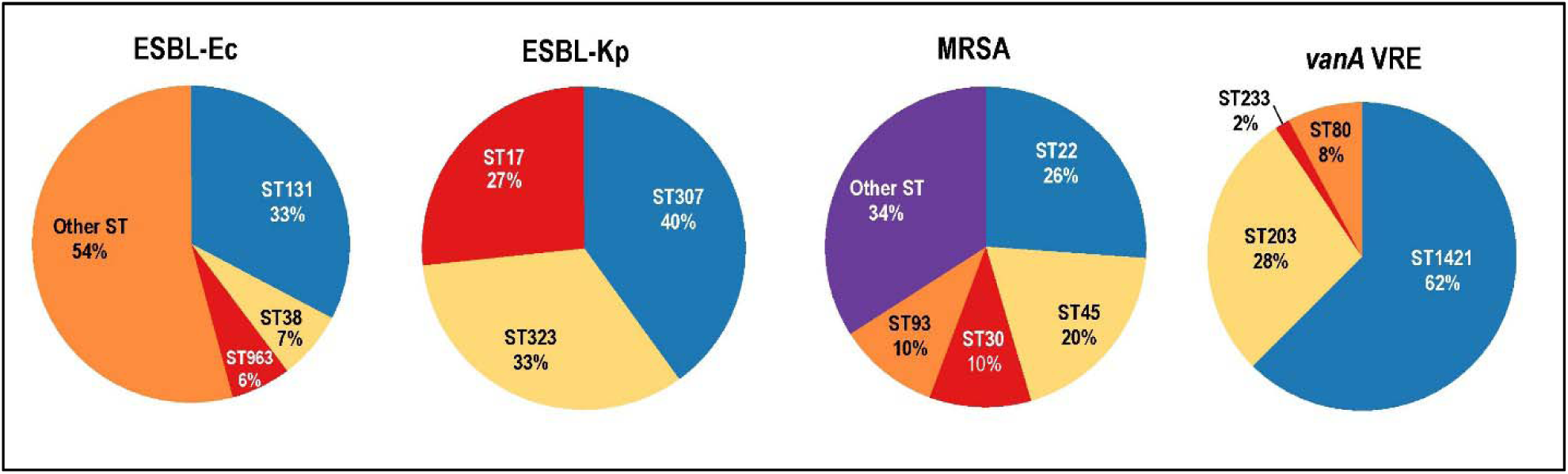
Most common multi-locus sequence types identified in this study. Abbreviations: ESBL-Ec, extended-spectrum beta-lactamase phenotype *E. coli*; MRSA, methicillin-resistant *S. aureus*; *vanA* VRE, *vanA*-producing vancomycin-resistant *E. faecium*; ESBL-Kp, extended-spectrum beta-lactamase phenotype *K. pneumoniae*; ST, sequence type.

#### MRSA

ST22 was most common amongst MRSA (25.6%), followed by ST45 (18.9%)(Figure 6). Some clones were more likely isolated in the first two days of admission (ST30 88.9% and ST93 100% isolated in first two days) compared with others (ST22 47.8% and ST45 58.8% isolated in first two days). Very few ST239 were isolated (dominant clone in several other Australian states).

#### ESBL-Ec

ST131 was by far the most common ESBL-Ec sequence type (71/218 isolates, 32.6%), followed by ST38 (6.9%), 963 (6.0%) and ST10 (5.0%)(Figure 6). Sequence types were otherwise quite diverse, with 45/56 STs having ≤2 isolates. CTX-M was the dominant ESBL gene group (85.5% of isolates), with CTX-M-15 being most common (43.0%). Two isolates had unexpected AMR genes detected (*mcr-1* and *rmtB*), prompting the implementation of enhanced infection control measures.

#### ESBL-Kp

ESBL-Kp were polyclonal, with only three STs represented by more than one isolate (Figure 6). CTX-M-type ESBL genes were also the most common in ESBL-Kp (87.1%).

### High rates of transmission detected for some MDROs

To investigate the potential transmission rates for major MDROs in this study, genomic comparisons were performed and genomic links to other study patients (pairwise SNPs at or below transmission screening threshold [see Methods]) were determined. Overall, 113/358 patients (31.6%) had potential genomic links to other study patients: 95.0% of *vanA* VRE, 23.3% of ESBL-Kp, 20.2% of ESBL-Ec and 11.6% of MRSA. Of these potential genomic links, 78/113 patients (69.0% under genomic link screening threshold) had probable transmission by epidemiology (see definitions in Methods), and a further 19 patients had possible transmission by epidemiology (Table 5).

**Table 5.** Likelihood of MDRO transmission by epidemiology by species.

| Likelihood of transmission<br>by epidemiology<sup>a</sup> | Species |  |  |  | Overall<sup>b</sup> (%) |
| --- | --- | --- | --- | --- | --- |
|  | ESBL-Ec | MRSA | vanA VRE | ESBL-Kp |  |
| **Total no. of patients in**<br>**study with MDRO** | 203 (56.7%) | 86 (24.0%) | 60 (16.8%) | 30 (8.4%) | 358 |
| **No. of patients with**<br>**potential genomic links<sup>c</sup>** | 41 (20.2%) | 8 (9.3%) | 57 (95.0%) | 7 (23.3%) | 113 (31.6%) |
| **No. of patients (%) in each epidemiologic category** |  |  |  |  |  |
| Probable | 20 (9.9%) | 5 (5.8%) | 46 (76.7%) | 7 (23.3%) | 78 (21.8%) |
| Possible | 10 (4.9%) | 2 (2.3%) | 7 (11.7%) | - | 19 (5.3%) |
| Unlikely | 11 (5.4%) | 1 (1.2%) | 4 (6.7%) | - | 16 (4.5%) |
| Same patient | 6 (3.0%) | 2 (2.3%) | - | - | 8 (2.2%) |
| **No. of transmission events per 100,000 OBDs<sup>d</sup>** |  |  |  |  |  |
| Probable | 15.9 | 4.0 | 36.5 | 5.6 | 61.9 |
| Probable + Possible | 23.8 | 5.6 | 50.0 | 5.6 | 77.0 |
| **Wards associated with probable transmissions<sup>e</sup>** |  |  |  |  |  |
| Intensive care | 27.6% | 5.9% | 12.5% | - | 23.3% |
| High-risk wards<sup>f</sup> | 8.6% | 5.9% | - | - | 7.5% |
| Other acute wards | 21.0% | 47.1% | 37.5% | 100% | 27.1% |
| Subacute care<sup>g</sup> | 2.9% | 29.4% | 25.0% | - | 7.5% |
| Day ward/operating<br>theatre | 3.8% | - | 12.5% | - | 3.8% |

<sup>a</sup> Definitions of likelihood of transmission by epidemiology: Probable, patients admitted to same ward at the same time; Possible, patients admitted to same hospital at same time, or same ward within 60 days (but without overlapping stays); Unlikely, all other patients outside these definitions; Same patient, isolates from same patient at different times. Reference: [30]
<sup>b</sup> Some patients represented under >1 species, hence totals may add to more than overall number of patients.
<sup>c</sup> Potential genomic links: Isolates analyzed for core genome single nucleotide polymorphisms (SNPs) by species and ST; isolate pairs with SNP distances below the transmission screening threshold ( $\leq 15$ SNPs (MRSA) or $\leq 25$ SNPs (other species), excluding same patient pairs) were designated as 'potential genomic links' for further epidemiologic investigation.
<sup>d</sup> No. of patients with both genomic and epidemiologic links to other patients in the study. OBDs, Occupied bed days - number of patients admitted overnight (excluding mental health and hospital-in-the-home services).
<sup>e</sup> For some patient pairs, admissions overlapped in multiple wards.
<sup>f</sup> High-risk wards, includes hematology, oncology, renal ward (including renal transplant), and liver transplant wards.
<sup>g</sup> Subacute care, includes aged care, rehabilitation, palliative care and spinal wards.

Genomic data have not previously been used to define transmission rates in the hospital setting (based on patient throughput), but the precision of this technology now makes this potentially possible. The highest proportion of transmissions occurred in *vanA* VRE (36.5 probable transmissions per 100,000 OBDs), followed by ESBL-Ec (15.9 probable transmissions per 100,000 OBDs), ESBL-Kp (5.6 per 100,000 OBDs) and MRSA (4.0 per 100,000 OBDs)(Figure 7). No transmission was found for *Pseudomonas* and *Acinetobacter* isolates. Probable transmission occurred mostly in intensive care and acute wards (Table 2). There was no clear threshold separating the pairwise SNP distributions for pairs designated as ‘probable’, ‘possible’ and ‘unlikely’ transmission (Figure 8).

**Figure 7.**
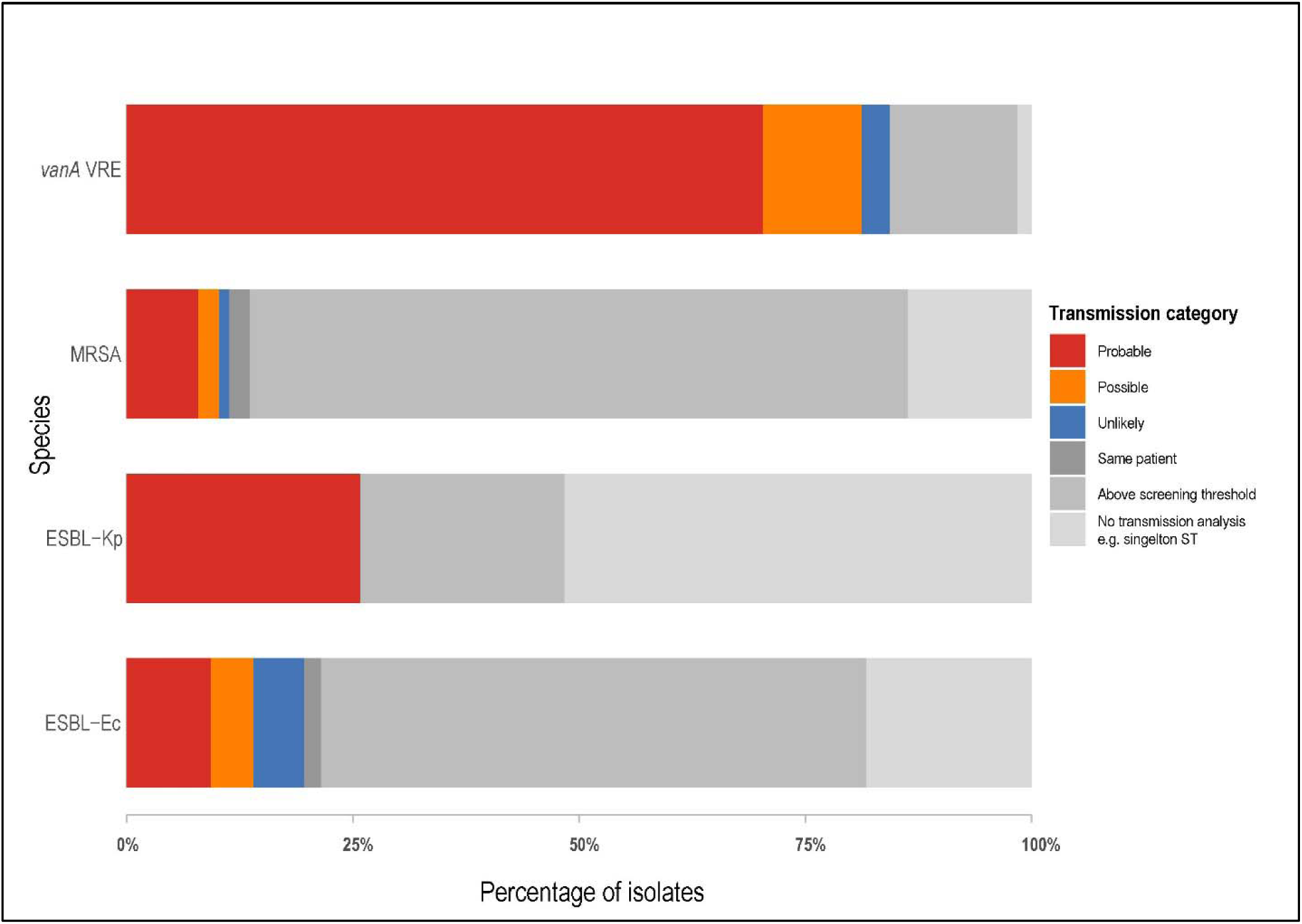
Transmission analysis results. **Transmission categories**: ‘Probable’, patients admitted to same ward at the same time; ‘Possible’, patients admitted to same hospital at same time, or same ward within 60 days (but without overlapping stays); ‘Unlikely’, all other patients outside these definitions; ‘Same patient’, isolates from same patient at different times; ‘Above screening threshold’, pairwise distances between isolates exceeded the transmission screening threshold (≥15 SNPs for MRSA, ≥25 SNPs for other species); ‘No transmission analysis’, isolates did not meet criteria for transmission analysis (ST only contained a single isolate, or only isolates from a single patient).

**Figure 8.**
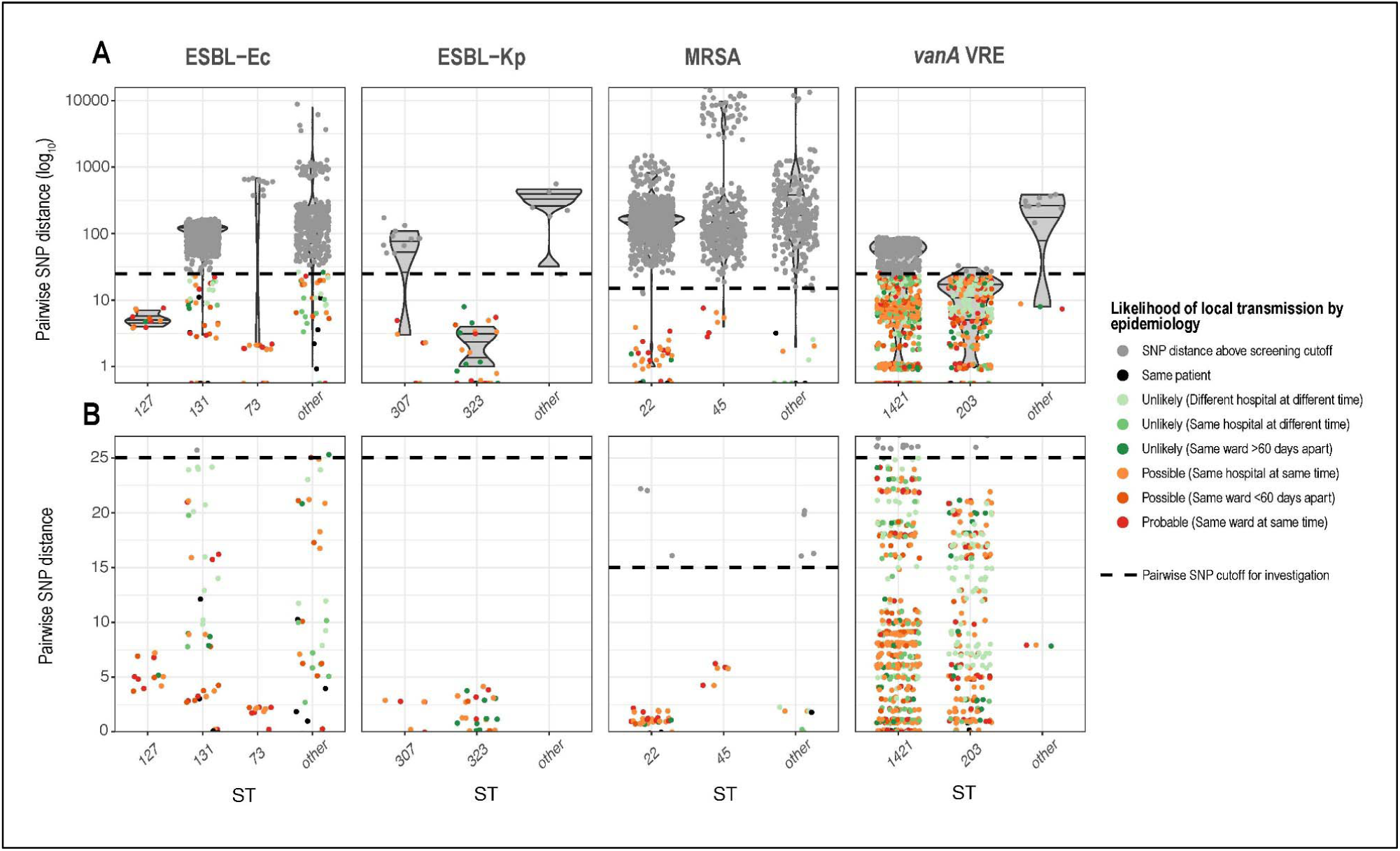
Transmission analysis: pairwise SNP distribution by species, ST and epidemiology. **Panel A:** overall view of pairwise SNP distances for each species, grouped by most common sequence types (ST), and other STs (note: log10 scale). **Panel B:** zoomed-in view of pairwise SNP distances for each species (linear scale). Dotted line represents transmission screening threshold of 15 SNPs for MRSA, and 25 SNPs for other species; bed move data only collected for patients with at least one isolate below this threshold. Each dot represents a pair of isolates; dots are colored by likelihood of local transmission by epidemiology (grey represents no data collected as pairwise SNP distance was above the transmission screening threshold).

## Discussion

In this study, we have collected comprehensive clinical and genomic data on four high-prevalence MDROs (ESBL-Ec, ESBL-Kp, MRSA and *vanA* VRE) and two low-prevalence, high-impact MDROs (CRPa and CRAb), established the local burden of infection and colonization with these MDROs, described the population structures, and used genomic data to infer putative MDRO transmission in hospitals, validated by epidemiologic data. ESBL-Ec was most common, with infection or colonization affecting at least 1.5 in every 1000 patients, higher than some other population-based estimates (e.g. 0.42 and 0.47 per 1000 patient-days in Canada 2009 and France 2013 [31, 32]). Similarly, the burden of MRSA in the study was relatively high, although still much lower than some estimates from other countries (e.g. 3 infections per 1000 patients [point-prevalence survey] in Canada in 2012 [33], compared to 0.42 per 1000 patient days in our study). Whilst *vanA* VRE was only moderately prevalent, this genotype only emerged in the last five years in Australia, where *vanB* continues to be dominant (21% of VRE *vanA* positive in a recent statewide survey, unpublished data). Interestingly, compared to other MDROs, ESBL-Kp was uncommon, indicating that this drug-resistant pathogen is not currently a major concern in our setting. Carbapenem-resistant *P. aeruginosa* and *A. baumannii* were infrequently isolated, this being a notable contrast to the high prevalence of these organisms reported in nearby Asian countries.

The combination of genomic surveillance and patient epidemiologic data revealed important information about which patients are affected by MDROs. Whilst the majority of MDROs were isolated within the first two days of admission (particularly for ESBL-Ec and MRSA), the time from admission to isolate collection had a biphasic distribution for most MDROs (i.e. many MDROs still isolated after the first week of admission, suggesting potential in-hospital MDRO acquisition), with the exception of *van A* VRE, where the majority of isolates were collected after the first week of admission. Interestingly, only 11% of patients in this study had an MDRO isolated without evidence of healthcare exposure (admitted from home, no admissions in last 12 months, and not known to be colonized with the same MDRO). Together, these data suggest that more of these MDROs are acquired in hospital than previously thought, challenging the prevailing dogma in the local infection control community (particularly for ESBL-Ec and MRSA).

*vanA* VRE exhibits several important differences compared to the other study MDROs; it was more likely to be responsible for colonization than to cause infection, more likely to be detected later in admission (suggesting hospital acquisition), more likely to cause infections in patients in high-risk wards, and had much higher rates of in-hospital transmission by both genomics and epidemiological classification (76.7%). Whilst there is ongoing debate about the pathogenicity of this MDRO [34], in our setting it is clearly a healthcare-acquired pathogen, especially in high-risk wards, and provides an opportunity for intervention, where genomics can be used to accurately target infection control interventions to wards with demonstrated transmission. Importantly, genomics can untangle complex transmission networks, identifying transmission in wards where patients were previously admitted (including general and subacute care wards, which together comprised over 60% of wards with probable *vanA* VRE transmission), rather than the ward on which the VRE was identified (often high-risk wards with routine screening, only comprised 12.5% of wards with probable transmission).

Comprehensive genomic epidemiology allows for analysis of MDROs on both a population level and at the level of individual patients. In describing the complex MDRO population structures, we observed both similarities and differences to other studies locally and internationally, such as the absence of ST1424 *vanA* VRE, dominant in adjacent states in Australia [13], and a lower proportion of ST131 ESBL-Ec compared to other studies internationally [35]. Genomic analysis also uncovered unexpected antimicrobial resistance genes *mcr-1* (encoding colistin resistance, a last-resort antibiotic which is not routinely tested in diagnostic laboratories) and *rmtB* (plasmid-borne AMR gene encoding broad-spectrum aminoglycoside resistance), prompting additional infection control measures for affected patients in this study, which would not otherwise have been detected, and could potentially modify antibiotic choices for these patients.

Genomic data has not previously been used to define an MDRO transmission rate based on patient throughput. However, this information could be potentially useful for benchmarking of hospitals, as well as potentially defining outcome measures in infection control intervention trials. Transmission analysis using a combination of genomics and epidemiology revealed a wide variation in the rates of transmission between different MDROs, with over 88% of *vanA* VRE isolates being designated as ‘probable’ or ‘possible transmission’ by study definitions. By contrast, the proportion of probable transmission for other MDROs was much lower (8.1% for MRSA to 23.3% for ESBL-Kp). Given that MRSA screening is uncommon in Victoria (as most hospitals do not isolate patients colonized with MRSA), the true rate of MRSA transmission is likely to be higher than seen in this study (as transmissions resulting only in colonization were not detected). Whilst the proportion of ESBL-Ec denoted ‘probable transmission’ was relatively low at 10%, the raw number of transmissions is significant (20 highly-related patient pairs) given how frequently ESBL-Ec is found in this population; patients infected or colonized by ESBL-Ec are not currently isolated in most hospitals in Victoria, due in part to the large numbers of colonized patients and scarcity of isolation rooms (see Table 4).

There are several limitations to our study, including variations in MDRO screening practices between participating sites, potential differences in collection of clinical isolates and microbiology workup between different hospitals, potential bias in recall and recording of hospitalization history in last 12 months (from patient recall and medical history review; no centralized database available), and absence of reliable data regarding patient overseas travel. Similarly, our transmission analyses may be limited by only being able to collect epidemiologic data (admission history, ward and bed moves) for patients with isolates below a screening threshold for genomic relatedness; this was chosen due to limited resources, as ward data was collected manually and hence quite resource-intensive. The bioinformatic methods used for transmission analysis are constantly evolving and not yet well-defined (multiple methods currently being used internationally), and limited in that they are only able to detect clonal transmission of whole MDRO bacteria, and are not yet geared to detect transmission of MDRO plasmids (due to limitations of short-read sequencing).

Despite these limitations, we believe that this study demonstrates the value of comprehensive genomic surveillance for MDROs on a population scale, a hospital scale and even at the level of the individual patient, and the potential for genomics to inform hospital infection control, if it is able to be applied in a timely manner. We plan to explore these concepts in a larger-scale translational study, using prospective genomics to detect transmission of hospital MDROs, in order to inform infection control interventions. Importantly, we need to be able to measure the potential benefits of genomics against the costs, in order to assess its likely utility in this setting.

## Acknowledgements

Other members of the Controlling Superbugs Study Group include:

Elizabeth Grabsch (Department of Microbiology, Austin Health), Joanna Price and Carolyn Tullett (Infection Control, Austin Health), Despina Kotsanas (Department of Microbiology, Monash Health), Louise Wright (Infection Control, Monash Health), Dr Suraya Hanim Abdullah Hashim and Jennifer Mitchell (Department of Infectious Diseases, Melbourne Health), Tram Nguyen and Margaret Savanyo (Department of Microbiology, Melbourne Health) and Peter Pham (MDU Public Health Laboratory, University of Melbourne).

The authors would like to gratefully acknowledge the assistance of the following people: Courtney Lane (statistics advice), John Greenough, Jennifer Breen and Penny Birchmore (assistance with infection control queries and patient movement data) and Carol Wedge (data entry).

